# How sample size influences the replicability of task-based fMRI

**DOI:** 10.1101/136259

**Authors:** Benjamin O. Turner, Erick J. Paul, Michael B. Miller, Aron K. Barbey

## Abstract

Despite a growing body of research suggesting that task-based functional magnetic resonance imaging (fMRI) studies often suffer from a lack of statistical power due to too-small samples, the proliferation of such underpowered studies continues unabated. Using large independent samples across eleven distinct tasks, we demonstrate the impact of sample size on replicability, assessed at different levels of analysis relevant to fMRI researchers. We find that the degree of replicability for typical sample sizes is modest and that sample sizes much larger than typical (e.g., N = 100) produce results that fall well short of perfectly replicable. Thus, our results join the existing line of work advocating for larger sample sizes. Moreover, because we test sample sizes over a fairly large range and use intuitive metrics of replicability, our hope is that our results are more understandable and convincing to researchers who may have found previous results advocating for larger samples inaccessible.

## Introduction

Recent years have seen an increased focus on the “reproducibility crisis” in science, both in science at large (Baker et al., 2016; Munafò et al., 2017; “Replication studies offer much more than technical details”, 2017), and perhaps even more acutely in the psychological sciences (Open Science Collaboration, 2015). Some of the reasons behind this crisis—including flawed statistical procedures, career incentive structures that emphasize rapid production of “splashy” (i.e., unlikely) results while punishing “failed” studies, and biases inherent in the publication system—have been articulated carefully in previous work, again both generally (Szucs, 2016; Barnes, Tobin, Johnston, MacKenzie, & Taglang, 2016; Wicherts et al., 2016), and for fMRI in particular (Carp, 2012; Button et al., 2013; Poldrack et al., 2017; Szucs & Ioannidis, 2017). Among these problems, the most frequently identified, and possibly the most easily remedied, is lack of statistical power due to too-small samples. Indeed, the field of fMRI has seen recommendations against large samples (e.g., Friston, 2012; cf. Ingre, 2013), and even when larger sample sizes are acknowledged as desirable, what constitutes “large enough” has often been an ad-hoc process of developing unempirical rules of thumb, or is based on outdated procedures (Thirion et al., 2007).

Of course, this lack of power is driven in large part by the great expense associated with collecting fMRI data (Mumford & Nichols, 2008). Even relatively small studies can cost several tens of thousands of dollars, and the funding system throughout much of the world is not generally set up to enable the routine collection of large (e.g., *N* > 100) samples. However, aside from these financial considerations, there are two other reasons researchers may persist in collecting small samples. The first is that while tools exist that allow researchers to do prospective power analyses for fMRI studies (Mumford & Nichols, 2008; Durnez et al., 2016), researchers may struggle to understand these tools, because defining power in an fMRI context involving hundreds of thousands of statistical tests is conceptually distant from defining power in a typical behavioral context, where there might be on the order of ten such tests. Relatedly, meaningfully defining “effect size” is conceptually straightforward in a behavioral context, but much less so in an fMRI context.

The second possible non-financial reason that researchers continue using small samples is because a number of studies have shown that fMRI has reasonably good test-retest reliability (Bennett & Miller, 2010; Gonzalez-Castillo & Talavage, 2011; Plichta et al., 2012; Bennett & Miller, 2013). It is possible that researchers take this to mean that large samples are not necessary, particularly if the researcher mis-understands standard design optimization approaches for increasing power at the individual level to mean their small samples are sufficiently powered (Liu, Frank, Wong, & Buxton, 2001; Wager & Nichols, 2003; Liu & Frank, 2004). However, test-retest reliability is not only not synonymous with replicability, but it is in some ways antithetical. This is because typical measures of test-retest reliability, e.g. the intra-class correlation (ICC), rely on variability across individuals. However, replicability is reduced by individual variability, particularly with small samples. While it is true that a measure with low test-retest reliability will have low replicability (in the limit, all individual maps are pure noise, and if there are suprathreshold voxels in the group average map, they likewise represent non-replicable noise), it does not follow that high test-retest reliability guarantees replicability at the level of group-average maps. Nor is it the case that variability between individuals in terms of brain activity is so minor that we can disregard it when considering the relationship between test-retest reliability and replicability; on the contrary, research has demonstrated that variability between individuals can swamp group-average task-related signal (Miller et al., 2009; Gabrieli, Ghosh, & Whitfield-Gabrieli, 2015; Dubois & Adolphs, 2016).

Our goal in the present study is to provide empirical estimates of fMRI’s replicability in terms of the levels of results that are useful in the field (i.e., multi-voxel patterns or cluster-based results, rather than, e.g., peak *t*-statistic values). Our specific focus is on the role of sample size (i.e., number of participants) on replicability, although we do examine the influence of other factors that might affect replicability, including design power (Turner & Miller, 2013). We also emphasize that our results, far from being relevant only to researchers whose specific interest is in studying reproducibility/replicability (e.g., Evans, 2017), are applicable to all researchers who are interested in using fMRI to produce valid and meaningful neuroscientific discoveries. In fact, we use *N ⋍* 30 as our standard for a “typical” fMRI sample size, which is in line with empirical estimates by Szucs and Ioannidis (2017) (median sample size of fMRI studies in 2015 = 28.5) and Poldrack et al. (2017) (75^th^ percentile of sample size in cognitive neuroscience journals published between 2011 – 2014 = 28). To preview our results, we provide an easily-interpretable demonstration of the facts laid out by Button et al. (2013) and Szucs and Ioannidis (2017): replicability at “typical” sample sizes is relatively modest, at best.

## Method

We carried out a series of analyses across eleven distinct tasks (from two large datasets). Because these analyses had the same form across all eleven tasks, we describe here the details of those analyses, and leave the description of the details specific to each task to the Supplemental Materials. We refer to the eleven tasks throughout this report using alphabetic labels A–K, because our interest is not in the identity of these tasks *per se*. We first describe the details of the first dataset (comprising tasks A–D), followed by a description of the second dataset (comprising tasks E–K), and end with a description of the analysis methods themselves.

### Dataset 1 (UIUC)

#### Participants

Participants were recruited from the Urbana-Champaign community as part of two separate intervention studies, each of which included a pre-intervention MRI session with two different fMRI tasks (for a total of four fMRI tasks). Both studies were approved by the University of Illinois Urbana-Champaign Institutional Review Board; all participants in both intervention experiments provided informed consent. All participants were right-handed, had normal or corrected-to-normal vision without color blindness, reported no previous neurological disorders, injuries, or surgeries, reported no medications affecting central nervous system function, were not pregnant, had no head injuries or loss of consciousness in the past two years, and were proficient in English. All participants received monetary compensation for participation. Only data provided at the pre-intervention time point (i.e., prior to the start of any intervention or experimental conditions) are included in the present analyses.

A total of 227 participants were recruited for and provided data in the first intervention study (Study 1). Task A includes a sample of 214 participants with complete data, and Task B includes 200 participants (of the 214 included in Task A) with complete data.

A total of 301 participants were recruited for and provided data in the second intervention study (Study 2). For the two fMRI tasks C and D, an identical set of 279 participants had complete data in both and are included in all analyses.

#### Scanning procedure

All participants in both Studies 1 and 2 were scanned on the same Siemens 3T Magnetom Trio. Study 1 participants were scanned with a 32-channel head coil; Study 2 participants were scanned with a 12-channel head coil. High resolution anatomical data were obtained using a high resolution 3D structural MPRAGE scan: 0.9 mm isotropic, TR = 1900 ms, TI = 900 ms, TE = 2.32 ms, with a GRAPPA acceleration factor of 2. Functional MRI BOLD data were collected using the Siemens echo-planar imaging sequence. Tasks B, C, and D used the following parameters: TR = 2000 ms, TE = 25 ms, flip angle = 90°, 92 *×*92 matrix with 2.5 mm in-plane resolution, 38 slices parallel to AC – PC with a 3.0 mm slice thickness and 10% slice gap. Task A used the same parameters, with the exception of the following: TR = 2360 ms, 45 slices with a 2.5 mm slice thickness. The number of repetitions varied for each task depending on the task duration (see Supplemental Materials for details). Finally, a gradient field map was collected for use in B0 unwarping matching the EPI parameters.

#### Preprocessing

Every run from each task was preprocessed identically using FSL’s (http://www.fmrib.ox.ac.uk/fsl) FEAT (FMRI Expert Analysis Tool, version 6.00) software package. Preprocessing included motion correction using MCFLIRT (Jenkinson, Bannister, Brady, & Smith, 2002), BET brain extraction (Smith, 2002), spatial smoothing with a 6 mm FWHM kernel, grand-mean intensity normalization, pre-whitening with the FILM tool (Jenkinson, Beckmann, Behrens, Woolrich, & Smith, 2012) and a high pass filter with a cutoff of (1/90) Hz. EPI images were additionally unwarped using the gradient field maps collected with the functional runs. The high-resolution structural scan was registered to the MNI152-T1-2mm standard brain via FLIRT (Jenkinson & Smith, 2001; Jenkinson et al., 2002) and further refined using the non-linear FNIRT tool (8mm warp resolution; Andersson, Jenkinson, Smith, et al., 2007). Transformation of each functional scan to the MNI standard brain was accomplished using a two-step process to improve alignment first by registering each EPI to the high-resolution structural scan with the FSL BBR tool (Greve & Fischl, 2009), and then applying the non-linear warp generated from the high-resolution scan to the functional scan.

#### GLM analysis

For a complete description of each task, task events, and contrasts, see Supplemental Materials. Briefly, Task A included 7 events; Task B included 4 events; Task C included 7 experimental events; Task D included 10 events. Predicted BOLD signals were generated for each event via convolution with a double gamma HRF (phase = 0). Six regressors derived from the motion parameters were included as regressors of no interest in each low-level model to mitigate the effects of motion in the data. The temporal derivative of each event was also included and the same temporal filtering that was applied to the preprocessed data was also applied to the model. A primary contrast of interest was identified for each task, defined by the cognitive effect that the task was designed to capture (i.e., the contrast an experimenter running any of these particular tasks would be primarily interested in). The contrast of interest was estimated in each subject in a mid-level analysis by combining all runs in a fixed-effects model. Following that, group-level statistical results for each task/contrast were generated using a mixed-effects model via FSL’s FLAME1 tool (Woolrich, 2008).

### Dataset 2 (HCP)

In addition to the data collected at University of Illinois Urbana-Champaign, we incorporated data from the Human Connectome Project (Van Essen et al., 2013; Barch et al., 2013). Details of the collection, preprocessing, and low- and mid-level GLM analysis of these data can be found elsewhere (e.g., the HCP S500 Release Reference Manual: http://www.humanconnectome.org/study/hcpyoungadult/document/500-subjects-data-release/). The volumetric analysis results (smoothed with a 4mm kernel) were downloaded from the Amazon Web Services S3 bucket designated for sharing these data (s3:// hcp-openaccess) in August 2017. A total of 463 participants from this release were included in our analyses (see Supplemental Materials for a full list of participant IDs); the remaining participants who were included in this release but not in our analyses were excluded on the basis of QC recommendations from the HCP group or due to errors encountered during analysis. Any participant who was excluded for a single task was excluded across all tasks, such that the seven HCP tasks have an identical set of participants. More details about the tasks and the contrasts we chose are available in the Supplemental Materials.

### Pseudo-replicate analysis

To estimate the replicability of group-level results, we took the following approach. First, we split our full sample of *N* participants into two randomized, non-overlapping sets (“P” and “Q”) of length *N/*2. Next, we chose a sample size *k* ∊ {16, 25, 36, 49, 64, 81, 100(, 121)} for which we sought to estimate the replicability, and used FSL’s FLAME1 tool to generate group-level statistical maps using the first *k* participants in both groups P and Q. Then, for each of a number of similarity measures, we computed the similarity between the P and Q group-level maps. Finally, we repeated the preceding steps across all in-range values of *k*, and for 500 random sorts in groups P and Q.

This same process was carried out for every task; for Tasks A and B, all sorts were done independently, while for Tasks C and D (which comprised an identical set of participants), the same 500 sorts were applied to both tasks, and likewise, a single set of 500 sorts was applied to Tasks E – K. For the purposes of presentation, we show the average replicability estimate across all eleven tasks for each sample size, along with the average within-task (and within-sample size) standard deviation, though we also include the curves for each task. Although we present error bars for all of our analyses, note that, as with all resampling-based analysis methods, our results suffer from complex interdependence that makes it difficult to draw strong inferences about differences between tasks. That is, the variance among the 500 simulated replications of a given task in our approach may underestimate the variance that would be observed given 500 true, completely independent replications of the task. Moreover, there is no analytic solution that would let us correct for this underestimation, if in fact it exists. Therefore, all error bars should be interpreted as being qualitative or illustrative, rather than as guides for whether differences are significant. To that end, we use standard deviations rather than standard errors or confidence intervals in our presentation of the results.

### Similarity statistics

The similarity statistics that we used to operationalize replicability were chosen to reflect different levels of focus. Broadly, there were three levels, which from most to least granular were voxel, cluster, and peak. We describe the measure(s) associated with each level in turn below. Throughout our analyses, we present results in an “exact replication” frame—that is, our results provide an empirical demonstration of what a researcher could expect if she were to re-run a study exactly, down to the sample size of the original study. Our gold standard would be to present results that reflect how well a study’s results capture “ground truth” as a function of sample size. Unfortunately, as is generally the case, the ground truth for the experimental contrasts we have included here is unknown.

Previous investigations in a similar vein have used either a meta-analytic approach or results from “large-enough” samples to approximate ground truth. However, meta-analyses suffer from well-established biases against small (but putatively nonetheless significant) results, and are moreover ill-suited to address some of the levels we focus on here. Likewise, to preview our results, although we have access to large samples by the standards of many neuroimaging studies, they may not be large enough to establish a reliable ground truth. More to the point, because of differences between tasks in terms of power and maximum available sample sizes, these ground truth maps would reflect different levels of “truthiness” across tasks, which would further confuse interpretation of these results. However, we do use results from the full sample in our “voxel-level (thresholded)” analyses, as described in more detail below.

#### Voxel-level replicability (intensity)

Arguably, the goal in fMRI is to accurately capture the activity in every single voxel. Indeed, many analysis methods are predicated on just such an assumption (e.g., MVPA, RSA, or encoding models), and many techniques for improving data acquisition or preprocessing are aimed at getting ever-finer spatial resolution (which we presume would be wasted effort if researchers’ goal was merely to approximate the spatial location of activity, or equivalently, the activity associated with a given location). Therefore, the first level of replicability on which we focused was the replicability of voxelwise intensities.

To quantify similarity, we used the Pearson correlation, which ranges from *-* 1 (inverse SPMs, invariant to scale) to 1 (identical SPMs, invariant to scale). The Pearson correlation gives us a holistic indication of how similar the between-voxel patterns of activity are across SPMs. To generate this measure, we computed the similarity between the vectorized unthresholded group-level SPMs, after applying a common mask to remove voxels that were zero (i.e., outside of the group FOV) in either SPM.

The null distributions for both metrics were constructed by generating SPMs of white noise spatially smoothed to match the observed smoothness in our real SPMs, and rescaled to equate the robust min and max (i.e., 2^nd^ and 98^th^ percentile, respectively). For each task and sample size, we generated the observed histogram of estimated FWHMs (using FSL’s `smoothest` command) as well as observed histograms of robust mins and maxes. We then parameterized these histograms and drew 1000 samples from the resulting parametric normal distributions. Finally, we generated 1000 maps of pure 𝒩 (0, 1) noise, smoothed each map with the corresponding sampled FWHM (using FSL’s `fslmaths` utilities) and rescaled to match the sampled robust min and max. We then computed the correlation between each real map and all 1000 of these null maps, and took the 95th percentile across these 1000 correlation values as the null for that specific real map; repeating this procedure across all maps yielded null values for every map, over which we took the average to arrive at the null curves presented in the figures.

#### Voxel-level replicability (thresholded)

Without abandoning the notion of describing replicability at the voxel level, it is nonetheless possible to relax the definition of what is being replicated somewhat—i.e., from raw intensity value to a binary “active”/“inactive” classification. To this end, we carried out a second set of analyses at the voxel level, using thresholded, binarized maps. As alluded to earlier, we used fullsample results in these analyses. Specifically, we thresholded each of the full-sample SPMs at liberal and conservative thresholds using FSL’s cluster-based thresholding (see next section for additional details on cluster-based thresholding), and used these thresholded maps in order to estimate the “true” proportion of voxels that should be suprathreshold for each task. Note that because the full samples comprise a larger number of participants than most fMRI studies—and because we will treat these full-sample results as “ground truth” which should be free of false positives inasmuch as possible—we have set the liberal and conservative thresholds higher than is typical in most fMRI experiments. The exact thresholds varied across data sets (in order to equate the power of the *z*-critical value as a function of sample size). See Table 1 for all *z* thresholds; all cluster *p* thresholds were set at *p* < 0.01.

**Table 1.** Specific z thresholds used in cluster-based thresholding of full-sample analyses for use in proportion-based thresholding analysis.

| Task(s) | Sample size | Liberal threshold | Conservative threshold |
| --- | --- | --- | --- |
| A | 214 | $\pm 3.50$ | $\pm 5.25$ |
| B | 200 | $\pm 3.39$ | $\pm 5.08$ |
| C–D | 279 | $\pm 4.00$ | $\pm 6.00$ |
| E–K | 463 | $\pm 5.15$ | $\pm 7.73$ |

With these per-task proportions suprathreshold (which are listed for each task in the Supplemental Materials), we simply applied proportion-based thresholding of the group-level SPMs (two-tailed) in order to match the full sample proportion suprathreshold. Conceptually, this is distinct from the cluster-based thresholding used in the subsequent sections, in that the voxels which end up suprathreshold are not guaranteed to meet any particular cutoff for significance, either at the voxel level or familywise. Thus, the quantity that is held constant across group-level maps P and Q is not the theoretical Type I or II error rates of each map, but simply the number of suprathreshold voxels. Our metric of replicability for these thresholded maps was the Jaccard statistic, which is simply the ratio of the intersection of a pair of thresholded maps divided by their union (with intersection calculated subject to the constraint that the voxel have the same sign in both maps—i.e., a voxel which was positive suprathreshold in P and negative suprathreshold in Q would not count as an intersection, but this voxel would be counted in the denominator). This statistic ranges from 0 (no overlap) to 1 (perfect overlap).

The null results were generated using the same approach outlined for the intensity-based voxel-level analyses, with the added steps of thresholding (two-tailed) the null maps at the same target proportion and computing the Jaccard overlap (again, in a sign-sensitive manner) between the pair of one null and one true map. As above, this procedure was repeated 1000 times and the 95^th^ percentile was taken.

#### Cluster-level replicability

While the ultimate or idealized goal of fMRI would seem to be voxel-level replicability, the common currency of today’s analytic landscape is generally the cluster (or as a special case, the peak; see next section). Therefore, the second level of replicability on which we focused was at the cluster level. Here, we chose to focus simply on the binary distinction between sub- and supra-threshold that forms the basis of cluster-based approaches (along with others). Although cluster-based approaches are widely used, it is less clear exactly what it means to replicate a cluster. Existing methods for conducting inferential statistics on clusters (e.g., Gaussian random field theory, Worsley, Taylor, Tomaiuolo, & Lerch, 2004; or permutation, Nichols & Holmes, 2002) refer to the null probability of observing a cluster of a given size (or possibly mass; Zhang, Nichols, & Johnson, 2009) conditioned on an initial threshold level, but do not address the question of exactly *where* this cluster appears.

Certainly, the spatial resolution at the cluster-level is coarser than at the voxel-level—researchers generally do not expect that every single supra-threshold voxel in a given cluster would be supra-threshold under replication, and likewise with sub-threshold voxels. Durnez, Moerkerke, and Nichols (2014), from which we take inspiration for our peak-based approach, employed a liberal definition in their cluster-based methods: a cluster is “replicated” if a single voxel from a given cluster is supra-threshold in replication. For our application, such a definition is far too generous, so we once again used Jaccard overlap. To generate clusters, we used FSL’s cluster-based thresholding on every group-level SPM, once at a liberal threshold (*z >* 1.96, *p <* 0.05) and once at a more conservative threshold (*z >* 2.81, *p <* 0.01). As with the previous thresholding analysis, we carried these analyses out in a two-tailed fashion, running FSL’s cluster-based thresholding once with *z > z*_*crit*_ and once with *z < − z*_*crit*_; then, Jaccard overlap was computed in a sign-sensitive manner.

We note as well some researchers might view cluster replication as a question of proximity; although Jaccard overlap is not a measure of proximity, it will generally track with proximity (i.e., as two clusters get closer together, their Jaccard overlap will increase). The exception to this is in the case of clusters which have zero intersection; a proximity-based measure would distinguish between a proximal pair of (nonintersecting) clusters and a distal pair, while both would have a Jaccard overlap of 0. In the interest of simplicity, as well as conceptual rigor when it comes to defining replication, we eschew such proximity-based measures.

The null was computed almost identically to that described in the previous section: each null smoothed map was thresholded (two-tailed) to match the proportion of supra-threshold voxels from the corresponding true image, and the Jaccard overlap between the two was computed.

#### Peaklevel replicability

The last analysis we report focuses on the level of peaks. Although clusters form the foundation of the majority of thresholding-based analyses used today, these clusters are typically reported simply in terms of the location and intensity of their peaks. In fact, some recent work has developed the statistical framework for understanding the behaviors of peaks, and how this can be used in, e.g., power analyses (Mumford & Nichols, 2008; Durnez et al., 2014). For the present purposes, we do not need to know the distributional characteristics of peaks, nor do we need to use the sophisticated estimation procedures described by Durnez et al. (2014). Therefore, we use the same cluster-extent (Gaussian random field theory) thresholding approach as for the cluster-level analyses. That is, whereas Durnez et al. (2014) use a peak-based secondary threshold when considering peaks as topological features, we use an extent-based secondary threshold.

A peak is considered replicated if it is suprathreshold under replication (i.e., part of any surviving cluster). This is a fairly generous definition of replication, but much less so than their cluster-level approach (i.e., non-zero overlap between clusters). Although we cannot interpret results in terms of false positives (because we are not comparing against “ground truth”), we can nonetheless examine the replication success of suprathreshold peaks. (We additionally provide an analysis of the relationship between peak *z* values and replicability for surviving peaks in the Supplemental Materials.) That is, we compute the proportion of peaks in one map that are suprathreshold in the complementary map. And unlike all previous measures, this measure is asymmetric—the proportion of P peaks that are suprathreshold in Q need not be equal to the proportion of Q peaks suprathreshold in P—so we calculated it in both directions and then averaged the results to arrive at the final value.

As with all other thresholding-based measures, we carried this analysis out in a sign-sensitive manner—i.e., a peak from a positive cluster did not count as overlapping. We used the same approach described in the preceding section to generate the null distribution. That is, we used the smoothed null maps, thresholded (two-tailed) to match proportion, to classify peaks.

### Measurables

Our expectation was that sample size would be the largest driver of replicability, irrespective of how it was measured. However, we also expected variability between our tasks (which would be unexplainable by *k*), as well as variability within a task for a given *k* (which would be unexplainable both by *k* and by task-level variables). Therefore, we carried out an analysis in an attempt to find other easily-measured variables that might explain these two types of variability. Although our primary goal is descriptive—that is, to identify the relationships present in our data—we used a modeling approach that in principle should allow generalization.

Before we describe this approach in detail, we note that we cannot use standard regression techniques to derive inferential statistics for our regressors, because our observations are non-independent (i.e., the correlation between any P and Q group-level maps reflect contributions from specific participants, all of whom will almost surely be members of other P or Q groups). Moreover, the influence of this non-independence varies across sample sizes, because the average number of participants in common between any two groups across iterations at a sample size of, say, 16, will be much lower than the average number of participants in common between groups at a sample size of 100.

Our modeling approach was relatively straightforward. First, we calculated per-pseudoreplicate measures for each of five variables: motion (taken as the average across functional runs of FSL’s estimated mean absolute RMS per run); contrast power (average across functional runs of the reciprocal of the contrast precision = c*inv(X’*X)*c’ where c is the contrast vector and X is the convolved design matrix); the number of outliers in the design matrixÕs hat matrix (average across functional runs of the count of diagonal entries on the hat matrix exceeding 2*rank(hat)/nrows(hat), where nrows(M) is the number of rows in M); within-individual variability (the average across every pair of runs of the whole-brain correlation between the run-level SPM for the given contrast within a participant) and between-individual variability (the whole-brain correlation between individual-level SPMs for the given contrast for each pair of participants).

The first four of these measures are defined on a per-participant basis, while the last is defined at the level of participant pairs. In order to translate these measures to the pseudoreplicate level, we took the arithmetic mean of the per-participant or -participant-pair measures over all participants in a given pseudoreplicate, as well as the standard deviation. Finally, to translate these per-pseudoreplicate measures to the pseudoreplicate-pair level (which is the level at which our outcome variables are defined), we took the arithmetic mean and absolute difference between the P and Q pseudoreplicate-level measures in each pair. Thus, each of our five original variables that varied at the individual level is expanded to four variables for the purposes of modeling. The final variable, sample size, does not vary below the highest level, and so does not need to be expanded similarly. Likewise, our outcome variable of unthresholded voxelwise similarity is already defined at the appropriate level.

Once we have the explanatory variables defined for every pseudoreplicate pair for every sample size and task, we did simple task-wise ordinary least squares regression to estimate the influence of each variable. First, for each task, we removed all observations from the largest sample size per task for the outcome variable and all explanatory variables (because for some tasks for which *k* was near *N*/2, some of our variables had variance near zero), then demeaned all variables. Next, we orthogonalized each of the four within-subjects variability regressors with respect to the twelve preceding variables (i.e., those derived from motion, contrast power, and hat-matrix outliers), and orthogonalized each of the four between-subjects variability regressors with respect to the preceding sixteen. Finally, we regressed the outcome variable on this set of explanatory variables (removing collinear variables as needed—this only occurred for tasks for which there was no variance in contrast power across individuals, such that some of the variables derived from contrast power were undefined).

We present two measures of the strength of the relationship between each original explanatory variable and the outcome variable. The first of these measures is the effect size of the largest of the four variables derived from each original variable—that is, 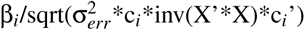, where 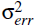 is the sum of squared residuals from the model, c_*i*_ is a vector of zeros with a ‘1’ as its i^*th*^ entry, and X is the design matrix. This is the standard formula for the *z*-value associated with a given β, without the reciprocal of the degrees of freedom in the denominator. The second measure is the increase in *R*^*2*^ that results from including adding all four variables derived from one of the original explanatory variables, compared against a model that includes all variables except these four.

In order to aggregate each of these measures across tasks, we computed the maximum a posteriori estimate of each across tasks. For the measure of effect size, we used a prior distribution of 𝒩 (0, 1) to shrink our estimated average across tasks. For the measure of Δ*R*^*2*^, which ranges from 0 – 1, we first used a logit transform to convert the measures to a scale with infinite support, then used a prior distribution of 𝒩 (−20, 10) in the transformed scale to shrink the estimate average toward −20 (again, in the transformed scale), and finally used the logistic transform to return the result back to the original 0 – 1 scale.

Note that this modeling approach is not able to probe the relationship between the explanatory and outcome variables at the level of differences between tasks. This was an intentional choice on our part because for some of the explanatory variables, between-task differences were many orders of magnitude larger than within-task differences. Moreover, because our tasks differ in many ways that aren’t captured by our chosen variables, it would be extremely speculative to attribute between-task differences in replicability (with a sample of only 11 tasks) based on a model including over twenty regressors. Therefore, using an approach that relies on the consistency of the within-task relationship across tasks is conservative, although we still caution readers against drawing strong inferences from our results, because our approach is designed to be primarily descriptive.

## Results

We present the result from each of the three levels of analysis described in the Methods—voxel, cluster, and peak—in separate sections below. Each section includes the true observed results for the measure used at that level in terms of the impact of sample size and task on that measure, as well as null results. In a separate section, we explore the relationship between various measurable properties of the data and the voxel-level replicability results.

For all Figures throughout the first three sections below, note that we plot results as lines for clarity, but computed our measures only for the discrete sample sizes marked on each *x*axis. Note too that the *x*axis uses a compressive (square root) scale.

### Voxel-level results

Our first analysis assessed the replicability of voxelwise patterns of raw SPM (“statistical parametric map,” in this case representing parameter estimates from the GLM; not to be confused with the software package of the same name) values, which we measured using Pearson correlation of un-thresholded P and Q maps. The results of this analysis are shown in Figure 1, which illustrates the results for the average across the eleven tasks, alongside the average of the null results across the tasks.

**Figure 1.**
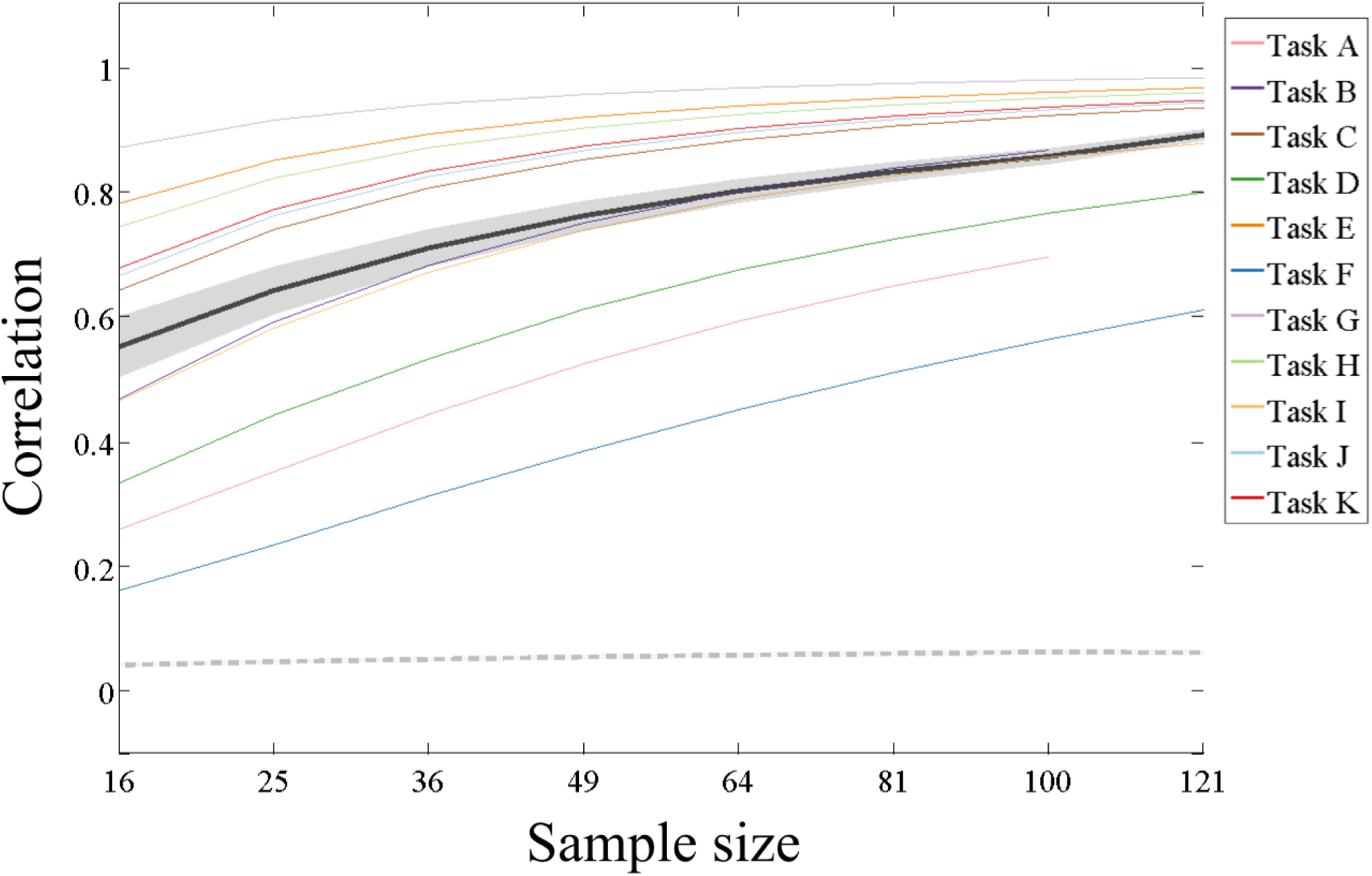
Replicability results for voxel-level (unthresholded) analyses. Average observed (±1 standard deviation) shown in black (dark gray); average null (± 1 standard deviation) shown in dashed medium gray (light gray). Also shown are individual task curves for Tasks A – K (colors given in legend).

There is no universally accepted value for this sort of replicability that would allow us to identify a minimum recommended sample size. However, we note that the smallest (measured) sample size for which the average *R*^*2*^ surpassed 0.5 was 36, which is still larger than our standard for a typical sample size.

The results of our second voxel-level analysis, of binary thresholded SPM replicability (using Jaccard overlap of maps thresholded using a conservative threshold), are illustrated in Figure 2. Results using a liberal threshold are presented in Figure S1. For these maps, we thresholded to match the proportion of suprathreshold voxels to the observed proportion suprathreshold for each task’s thresholded fullsample analysis. That is, differences between tasks in terms of power lead to differences in terms of the proportion suprathreshold, which in turn largely explains the differences between tasks in these four curves. Even at a sample size of 121, the average Jaccard overlap across tasks fails to surpass 0.6.

**Figure 2.**
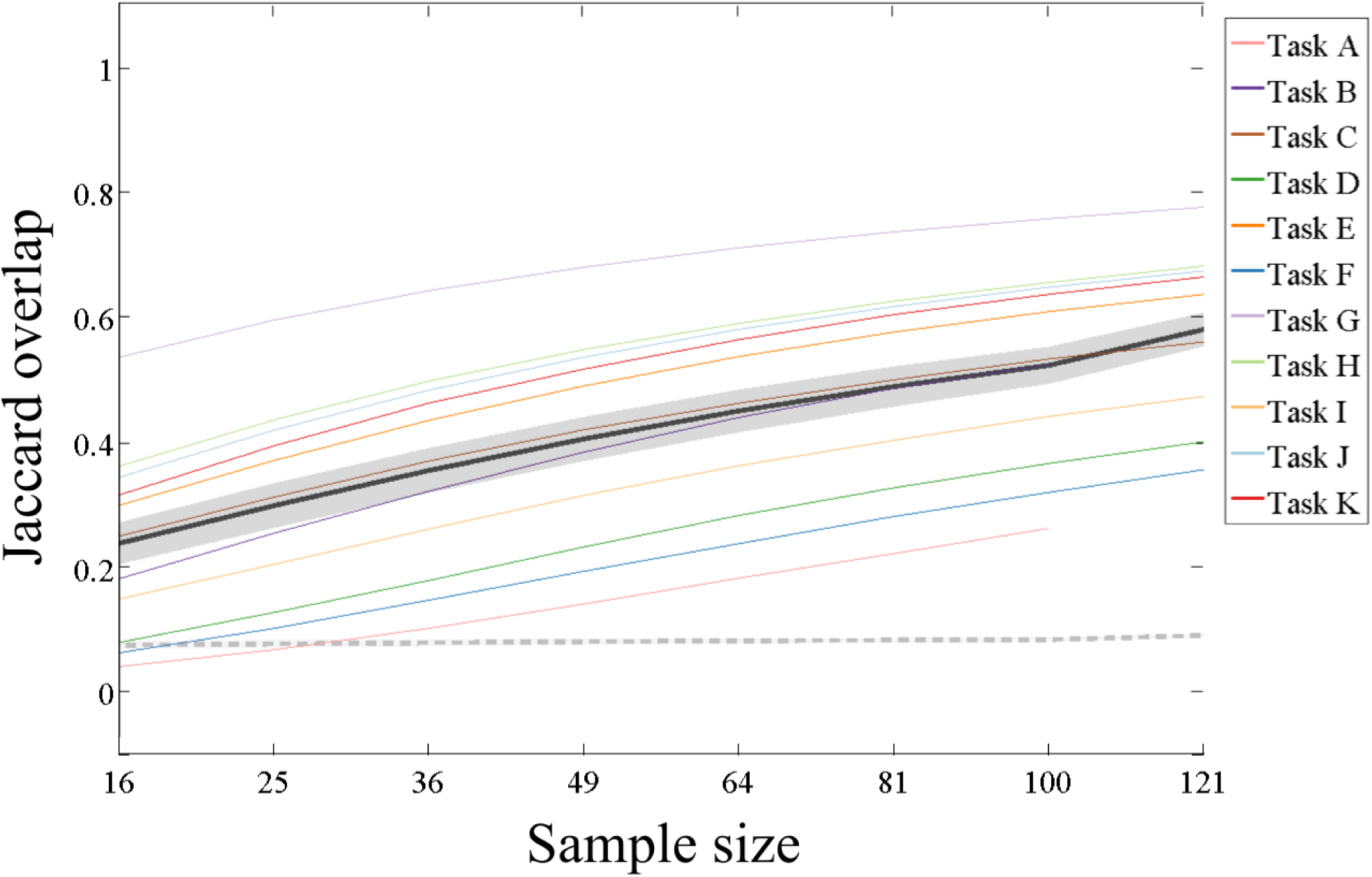
Replicability results for voxel-level (thresholded conservatively) analyses. See also Figure S1.

### Cluster-level results

The second level at which we considered replicability was at the cluster level. For this analysis, we thresholded each P and Q map using FSL’s cluster thresholding tool, and computed the Jaccard overlap between the resulting binarized thresholded maps. Figure 3 presents the results of our cluster-level analyses in terms of mean Jaccard overlap as a function of sample size for each task using the conservative threshold. Results using a liberal threshold are shown in Figure S2. Unsurprisingly, average Jaccard overlap at a sample size of 16 is near 0 for several tasks, because these SPMs are often null (i.e., contain no suprathreshold voxels), and even when both maps in a pair are non-null, the clusters overlap minimally. As with the analyses holding a set proportion suprathreshold per task, mean overlap remains below 0.5 up to a sample size of at least 81.

**Figure 3.**
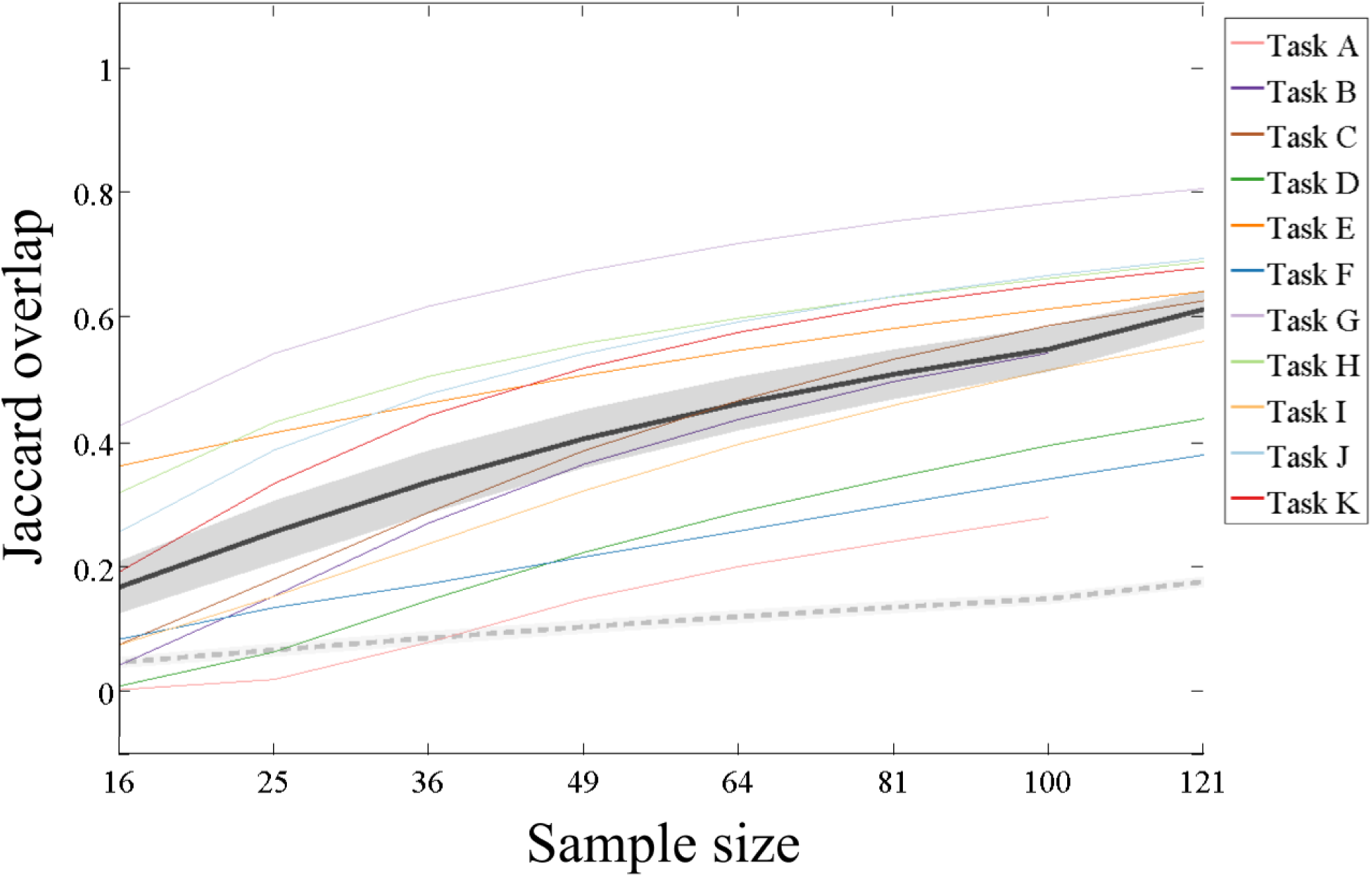
Replicability results for cluster-level (thresholded conservatively) analyses. See also Figure S2.

### Peak-level results

The final level of replicability we considered was at the level of cluster peaks. For this analysis, we assessed how frequently the peak voxel of each cluster was suprathreshold in its corresponding pseudo-replicate. We used a single peak per cluster (i.e., we ignored local maxima). Figure 4 illustrates the results for suprathreshold peaks. Results using liberal thresholds are shown in Figures S3. On average across tasks, even with a sample size of 121, the peak voxel failed to surpass threshold in its corresponding pseudoreplicate over 20% of the time.

**Figure 4.**
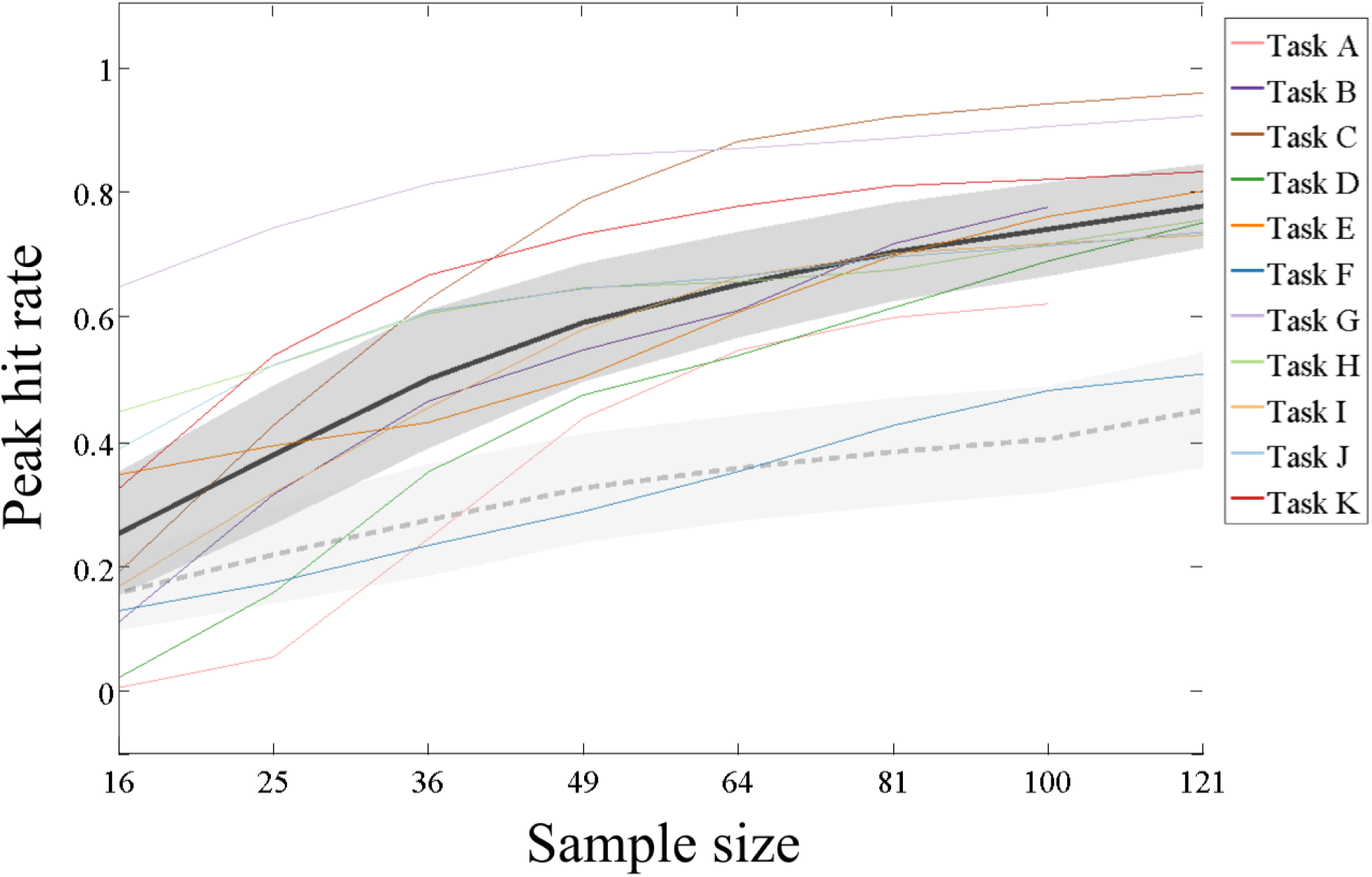
Replicability results for suprathreshold peaklevel (thresholded conservatively) analyses. See also Figure S3.

### Measurables results

Although our focus was on the effect of sample size on replicability, and this variable was the only one we manipulated systematically in our pseudoreplicate analysis, we nonetheless have access to a number of other variables whose potential influence on replicability we can measure, including variables related to motion, contrast power, and within- and between-individual variability. Our goal in these measurables analyses is not to draw strong inferences about the influence of each of these variables, which is complicated by the interdependence between many of our observations (e.g., overlap between pseudoreplicates in terms of which participants comprise each group; overlap between sample sizes for the same pseudoreplicate, with larger sample sizes constituting supersets of smaller sample sizes; and overlap between tasks in terms of participant identity). Instead, we present qualitative results of a simple analysis designed to provide some intuitive understanding of the relative role each variable plays in driving replicability. Full details on the modeling approach can be found in the Methods, but briefly, we fit a separate simple regression model relating each of the above-mentioned variables to replicability for each task, and take as our measures of interest the effect size and Δ*R*^*2*^ associated with each variable.

Only two variables emerged as qualitatively having more than a very weak relationship with replicability. As expected, sample size was related to replicability 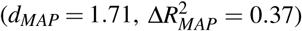 The second variable to demonstrate a relationship with replicability was between-individual variability 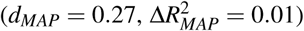. Although several other variables demonstrated a consistent relationship with replicability across tasks, the effects were minuscule. Results for all variables are given in the Supplementary Materials.

## Discussion

Despite the development of various tools meant to allow researchers to do prospective power analyses (Mumford & Nichols, 2008; Durnez et al., 2014), such tools are apparently used only infrequently by researchers. Several previous studies have suggested that neuroimaging studies suffer from a marked, possibly fatal, lack of statistical power (Button et al., 2013; Szucs & Ioannidis, 2017). However, Type II errors are not the only problem plaguing neuroimaging, as other studies have demonstrated that certain widely used falsepositive correction methods underestimate true false positive rates (Eklund, Nichols, & Knutsson, 2016), and multiple testing correction has been a topic of substantial investigation throughout the history of neuroimaging (Bennett, Miller, & Wolford, 2009).

This previous work has uncovered persistent and troubling problems with standard neuroimaging approaches, particularly as it regards the use of appropriately well-powered (i.e., large) samples. However, no prior work has operationalized replicability in the concrete, intuitive ways we have here, nor has any prior work systematically examined the impact of sample size and other dataset properties on such measures of the replicability of task-based fMRI. One exception is Thirion et al. (2007), which is a spiritual predecessor to the current work. However, in addition to being a decade old (during which time the field of fMRI research has changed substantially), the measures used by Thirion et al. (2007) may not be as intuitively accessible as those we have adopted here. Furthermore, that work focused on many other drivers of replicability (e.g., threshold choice, thresholding method), and their main conclusion with respect to sample size is that *N* = 20 should be considered a minimum—a value that our results suggest is too low.

Our results demonstrate that, regardless of whether one conceptualizes replicability as being about patterns at the level of voxels, clusters, or peaks, our estimates of replicability at typical sample sizes are quite modest. For instance, the mean peak hit rate across our eleven tasks for a sample size of 36 (which is above the mean or median sample sizes reported in Poldrack et al., 2017 and Szucs & Ioannidis, 2017 over the last several years)is below 0.5. The observed mean tells us that over 50% of the cluster peak voxels observed in a group SPM with this sample size will fail to be suprathreshold in an exact replication. Such a lack of replicability is likely to be concerning to researchers who rely on methods that assume a high degree of spatial specificity (e.g., MVPA). And this same pattern holds true, often to an even greater extent, across our other measures. Furthermore, our results represent a best case scenario for replicability (at any of our tested sample sizes for any of our tasks) because we drew samples from the same broad population, we collected all data at one site and one scanner, the experimental methodology and materials were exactly identical for all subjects, and all fMRI data processing was completed using identical processing pipelines on the same computers using the same software. In other words, for any single iteration in our bootstrap method, all pseudo-replicates could be classified as “exact” replications. Deviations from any of these criteria would likely introduce variability in the data collection and processing streams, yielding lower observed replicability.

What can explain this pattern of results? Clearly, there are two possible sources of noise in a group-average result: within-subject variance and between-subject variance. Increasing sample size reliably reduces the impact of both sources of noise. However, our analyses of how several easily-measured properties of each data set impacted replicability revealed small but consistent roles for several other factors, most notably the mean between-subject similarity. The idea of inter-individual consistency has been explored previously, and it is not altogether uncommon for researchers to publish maps demonstrating how consistent their results were across participants (e.g., Rosenblatt, Vink, & Benjamini, 2014; Seghier & Price, 2016). However, our reinforce the measurable impact of individual differences. A long line of research has highlighted this extraordinary variability, and argued for taking advantage of this variability, or at least acknowledging and attempting to control for it (Miller et al., 2002; Van Horn, Grafton, & Miller, 2008; Miller et al., 2009; Miller, Donovan, Bennett, Aminoff, & Mayer, 2012). Our results fit with these earlier observations that individual identity is a driver of patterns of brain activity. Moreover, to the degree that our scanned samples were more homogeneous than the population at large (as is generally the case of scanned samples that largely comprise undergraduates or members of the campus community), it is reasonable to expect that the influence of individual differences would be even larger in any study that used truly representative sampling.

It is possible that our results do a poor job of capturing the average replicability that should be expected across the field at large. However, we do not believe this to be the case, for four reasons. First, our results for our tasks C and D, as well as for tasks E – K, each of which separately included identical sets of participants and exactly matched pseudo-replicate groups, span a fairly wide range of replicability values. Second, the results from our two distinct datasets were broadly similar, despite myriad differences at all levels of data collection and analysis. Third, the included tasks are well-known, and cover a number of cognitive domains of general interest to researchers in cognitive neuroscience. And fourth, our results are consistent with earlier work demonstrating the inadequacy of “typical” (*N* ⋍ 30) sample sizes. Although there is no simple way to map our results onto these earlier studies, the general conclusion is much the same.

It is also possible that results using other methods—for instance, multivariate methods—would demonstrate substantially higher replicability than the results we present for the GLM. Although it is beyond the scope of the present work to adapt our approach to other analysis methods, part of our goal in using data from the Human Connectome Project (Van Essen et al., 2013) was to enable other researchers to carry out their own analyses of these data using a similar approach, making whatever changes they see fit (in preprocessing, software tool, or analysis method).

Although our results suggest that typical sample sizes are inadequate, it would be inappropriate for us to try to use our findings to identify a universal “minimum” sample size that could be adopted across the field. This is because our results do not represent how well sample sizes approximate “ground truth” but rather the expected replicability at each sample size. Moreover, although our tasks cover a reasonable range of effect sizes (as demonstrated by the different replicability estimates across tasks), any universal recommendation would have to be made for the smallest “meaningful” effect size, which is not an agreed-upon quantity in the field, and which is probably smaller than the smallest effect size we observed. Instead, we point readers to existing tools for conducting prospective power analyses, and hope that future research will develop similar tools that make use of the replicability measures we have employed here.

Our hope is that whereas earlier work pointing out the ubiquity of underpowered studies may have been seen by the average researcher as too abstract or technical to worry about, the present results are accessible enough that researchers can see that “typical” sample sizes produce only modestly replicable results, irrespective of how replicability is measured. Thus, our results add to the growing consensus calling for a shift in the field, away from small-scale studies of hyper-specific processes to large-scale studies designed to address multiple theoretical questions at once. Alternatively, methods which are transparent about treating individuals as unique—for instance, individual differences approaches (Van Horn et al., 2008) or encoding methods (Naselaris, Prenger, Kay, Oliver, & Gallant, 2009)—likely deserve more attention for their potential to overcome at least one part of the problem with small samples (i.e., individual variability).

## Conclusion

Replicability is the foundation of scientific progress. Unfortunately, for a variety of reasons, many scientific fields are currently gripped by a crisis of irreproducibility (Baker et al., 2016). While some of the causes of this crisis are deeply interwoven into the academic landscape—incentives related to publication, funding, and tenure—the most straightforward solution relates to statistical power. Researchers in fMRI may have believed that they were adequately addressing concerns about power by using carefully optimized designs and rule-of-thumb “large enough” sample sizes (Thirion et al., 2007; Friston, 2012; Liu et al., 2001). Indeed, the success of quantitative meta-analysis methods (e.g., activation likelihood estimation; Eickhoff, Bzdok, Laird, Kurth, & Fox, 2012), alongside reports of moderate test-retest reliability for task-based fMRI (Bennett & Miller, 2010), may have reinforced the sense that power in task-based fMRI was a solved problem. However, meta-analytic approaches work precisely by relaxing specificity about spatial location (and in many cases, about design features including task, contrast, or putative cognitive processes); likewise, test-retest reliability is only weakly related to replicability. Despite empirical work demonstrating that typical fMRI sample sizes are inadequate, there seems to be little motivation to change the status quo (Button et al., 2013; Szucs & Ioannidis, 2017). Our results unambiguously demonstrate that replicability (as measured at multiple levels of analysis) is strikingly low at “typical” sample sizes, thus serving to highlight and extend these previous results. The solution to this problem may be arduous for researchers and funding agencies, for instance requiring a paradigm shift away from incremental research using bespoke experiments with small samples. However, if our goal is the advancement of scientific understanding, the status quo—thousands of underpowered and minimally-replicable papers published annually—clearly cannot continue.

## Acknowledgments

The research is based upon work supported by the Office of the Director of National Intelligence (ODNI), Intelligence Advanced Research Projects Activity (IARPA), via Contract 2014-13121700004 to the University of Illinois at Urbana-Champaign (PI: Barbey). The views and conclusions contained herein are those of the authors and should not be interpreted as necessarily representing the official policies or endorsements, either expressed or implied, of the ODNI, IARPA, or the U.S. Government. The U.S. Government is authorized to reproduce and distribute reprints for Governmental purposes notwithstanding any copyright annotation thereon.

The authors would like to thank Soohyun Cho, Keith Holyoak, Michael Stevens, Jeremy Gray, Todd Braver, and Debbie Hannula for graciously providing original task design, timing, and stimulus files for the tasks reported in this manuscript. The authors also thank Jennifer Elam and Matthew Glasser for their help in retrieving the HCP data.

## Supplemental Materials

### Task descriptions (design and analysis)

The design of each task was based closely on a previously-published instantiation of each task. Here, we provide the basic details of each task, and explicitly highlight any points at which the design or analysis deviated from its previously-published antecedent.

**Task A.** See Gray, Chabris, and Braver (2003) for full details regarding the paradigm. This was a 3-back working memory task. Participants saw multiple short series of consecutive stimuli, during which they had to respond to items that had appeared exactly three items earlier (“targets”). These were intermixed with new items, as well as items that had appeared either two, four, or five items earlier (“lures”). As in Gray et al. (2003), there were two functional runs (one using faces, the other using words, order counterbalanced across participants), each of which included four blocks of 16 trials (plus five jitter fixation trials per block). Trials were modeled with seven regressors: two each (correct/incorrect) for targets, lures, and non-lures; and one for missed trials. Our primary contrast of interest compared correct targets and correct lures. On average per run, this contrast included 10.1 trials (standard deviation = 2.7 trials) versus 12.8 trials (standard deviation = 2.3 trials).

**Task B.** See Hannula and Ranganath (2008) for full details regarding the paradigm. This was a task of relational memory. Participants viewed displays of four 3D objects on a 3 *×* 3 grid, and had to indicate whether a test grid, displayed rotated after a short delay, matched the original layout. These test grids could be of three types: “match,” in which all items retained their original relative positions; “mismatch,” in which one item moved out of position; or “swap,” in which two items swapped positions. Each trial was comprised of an encoding period, a delay period, and a test period. There were five functional runs, each of which included 15 trials. These trials were modeled with a simplified set of four regressors: one each for correct encoding + delay periods (collapsed across trial types), match test periods, and non-match test periods (collapsing across “mismatch” and “swap” trials); and one for all periods of all incorrect trials. Our primary contrast of interest compared correct match and nonmatch test periods. On average per run, this contrast included 3.9 trials (standard deviation = 0.7 trials) versus 5.7 trials (standard deviation = 1.9 trials).

**Task C.** See Cho et al. (2010) for full details regarding the paradigm. This was a task of analogical reasoning, with a 2 × 2 design in which relational complexity (the number of tobeattended stimulus traits, 1 or 3) was crossed factorially with interference level (the number of irrelevant dimensions that lead to an incorrect response, 0 or 1). In our adaptation of their design, we included three functional runs, each of which contained 54 trials. These trials were modeled by seven (RT-duration) regressors: four defined per the 2× 2 design described above; another two for invalid trials (relational complexity 1 or 3); and a final regressor for error trials. Our primary contrast of interest compared relational complexity 1 with relational complexity 3, collapsing across interference levels. On average per run, this contrast included 18.5 trials (standard deviation across participants = 1.3 trials) versus 17.1 trials (standard deviation = 2.4 trials).

**Task D.** See Witt and Stevens (2013) for full details regarding the paradigm. This was a task of set switching. Participants were always tasked with counting the number of unique levels of a given relevant dimension; the relevant dimension changed (as indicated by a printed cue above the stimulus) every 1–6 trials. Trials varied in terms of: switch vs. non-switch (as well as number of preceding non-switch trials for switch trials); stimulus complexity (1, 2, or 3 varying dimensions with multiple levels); and response complexity (1, 2, or 3 potential valid response options across all dimensions). As in Witt and Stevens (2013), there were two functional runs, each with 81 trials. These trials were modeled with ten (RT-duration) regressors: two for switch/non-switch; six parametric regressors (orthogonalized with respect to the switch/non-switch EVs) encoding separately for switch and non-switch trials stimulus complexity, response complexity, and number of preceding non-switch trials; and two regressors to model error and posterror trials. Our primary contrast of interest compared switch and non-switch trials. On average per run, this contrast included 31.0 trials (standard deviation = 5.6 trials) versus 32.7 trials (standard deviation = 5.0 trials).

**Tasks E–K.** See Barch et al. (2013) for details on the tasks included in the Human Connectome Project. Table S1 lists the tasks according to the alphabetic identifier associated with each here, as well as the contrast chosen for each task. Table S3 lists the subject IDs for all 463 participants included in the present analyses. We additionally note that HCP’s core research team recommends caution in the use of volumetric data; however, because our aims are orthogonal to those of most users of these task-based data, and because our measures are all designed for volumetric data, we feel that our use of these data, rather than the surface-based data, is appropriate.

### Peak height replicability analysis

For our peak analyses, in contrast to our other analysis approaches, it is possible to construct a disaggregated statistic—that is, to define replicability on a peak-by-peak basis, rather than only mapwise. This allows us to look in a more fine-grained manner at the relationship between effect size, replicability, and sample size. To this end, we collated each peak *z* value with whether that voxel was replicated (i.e., was suprathreshold in the counterpart map), separately for each sample size but combining across all tasks. We then fit a separate logistic regression model for each sample size. We then used the `sim` function (part of the `arm` package in **R**) to graphically display uncertainty around the model fits, which is especially pronounced for values of peak *z* outside of the range observed for a given sample size. Note that this approach treats task as a fixed effect, and moreover, weights tasks proportional to the number of total peaks across all maps for a given sample size. Note too that, as with the main peak analysis reported in the manuscript, a low *p* (replicated) is heavily influenced by the sparsity of the counterpart map. The results of this analysis are shown in Figure S4.

## Supplemental figures

**Figure S1.**
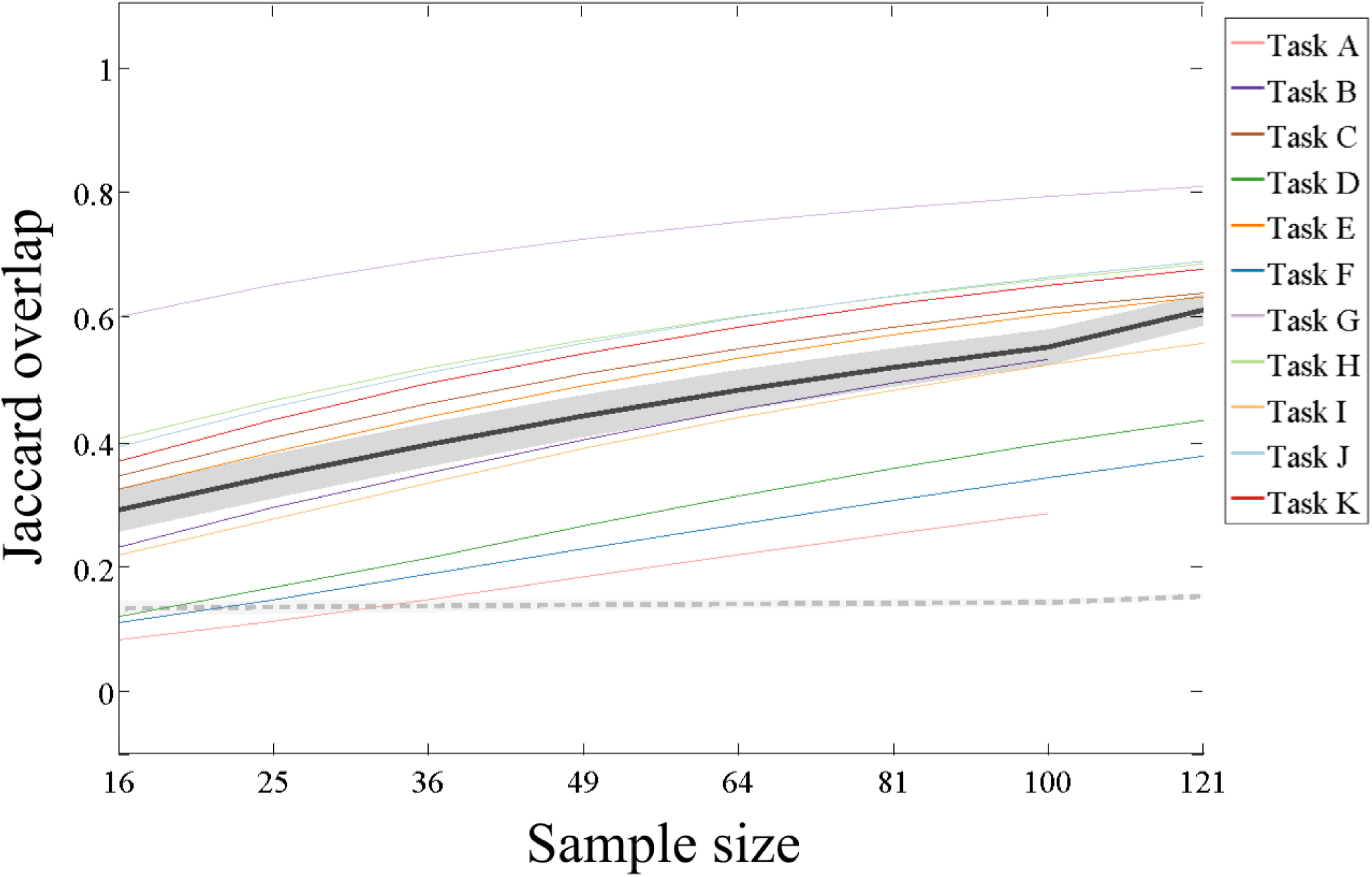
Replicability results for voxel-level (thresholded liberally) analyses.

**Figure S2.**
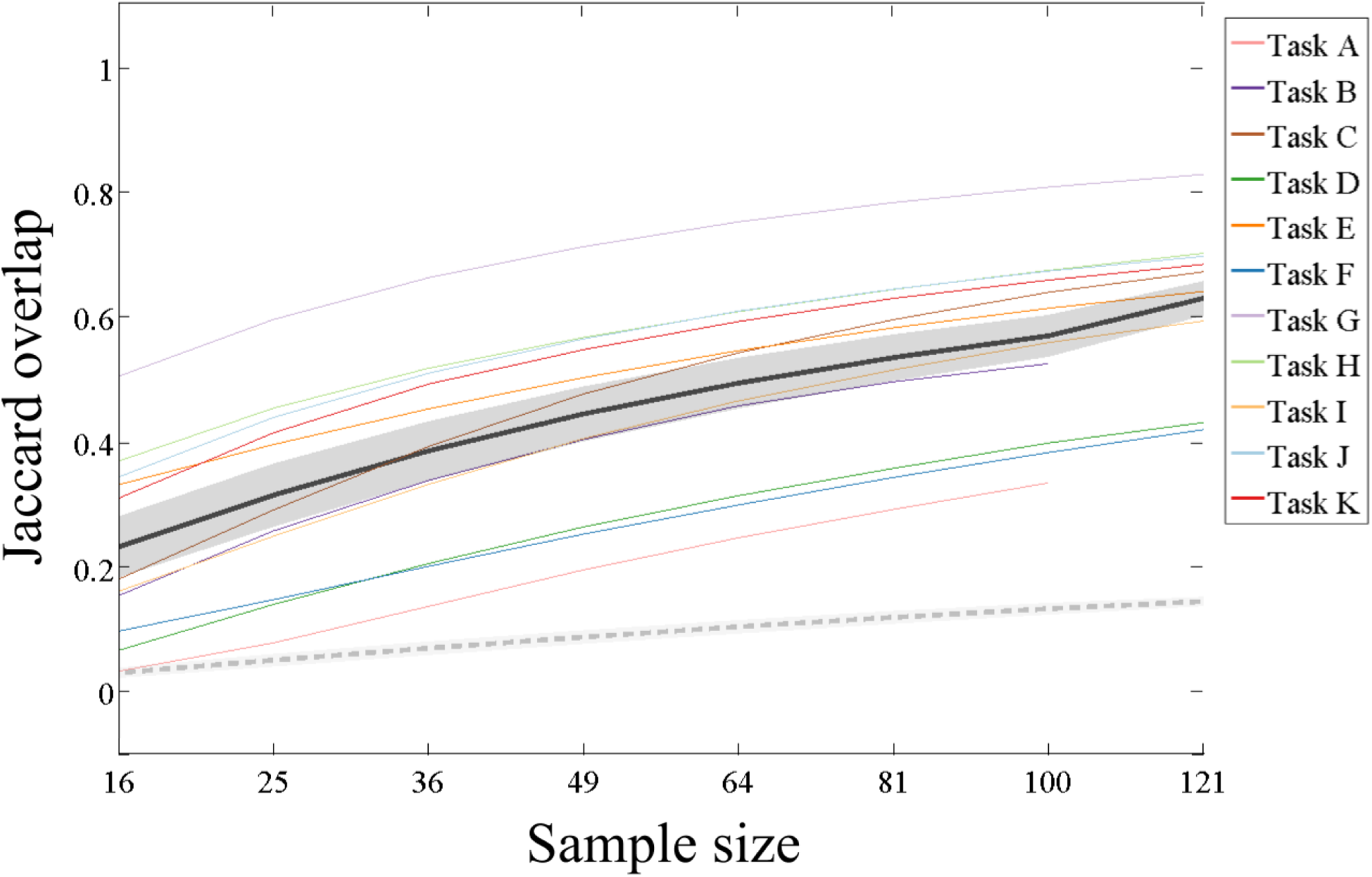
Replicability results for cluster-level (thresholded liberally) analyses.

**Figure S3.**
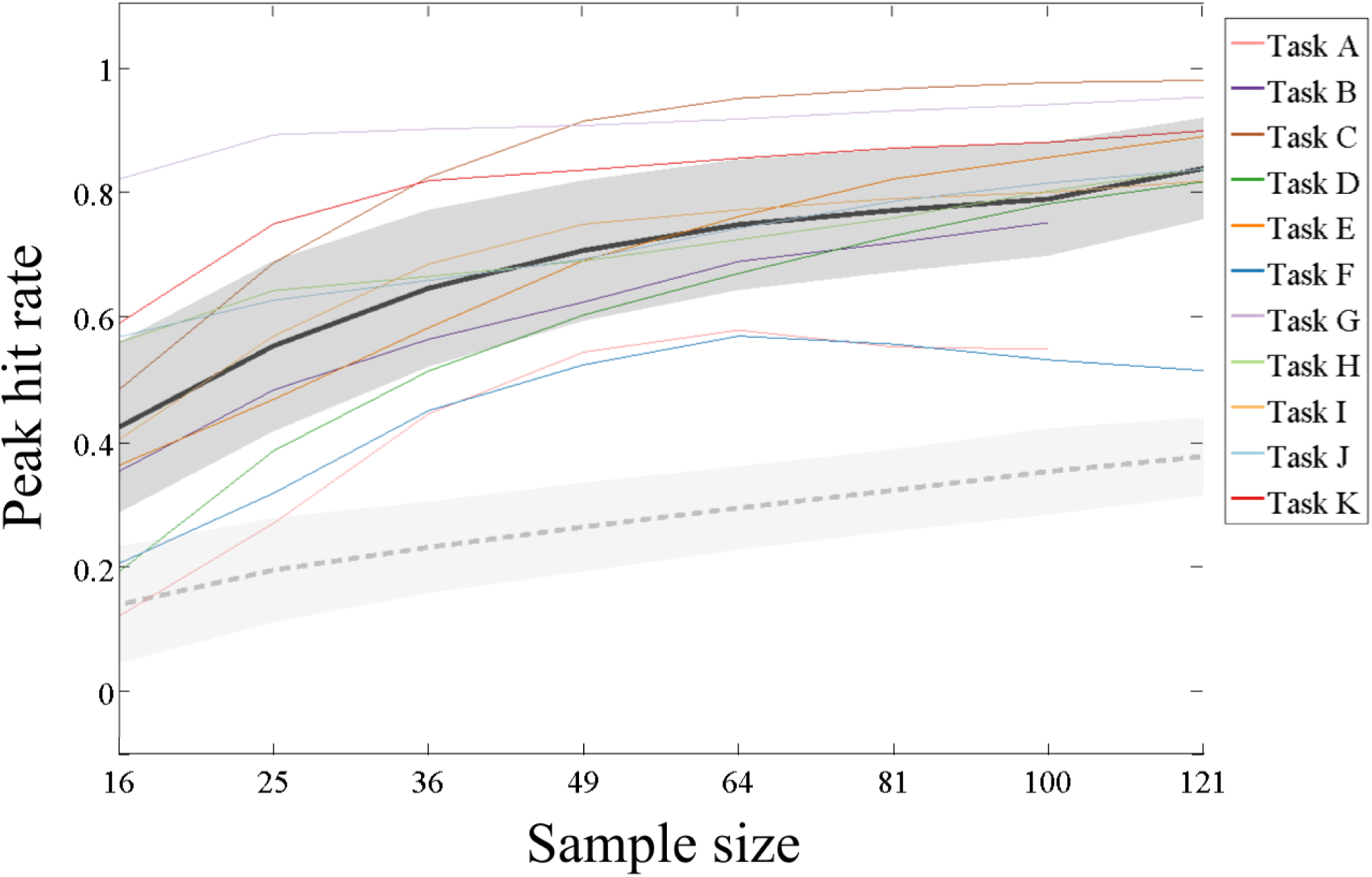
Replicability results for suprathreshold peak-level (thresholded liberally) analyses.

**Figure S4.**
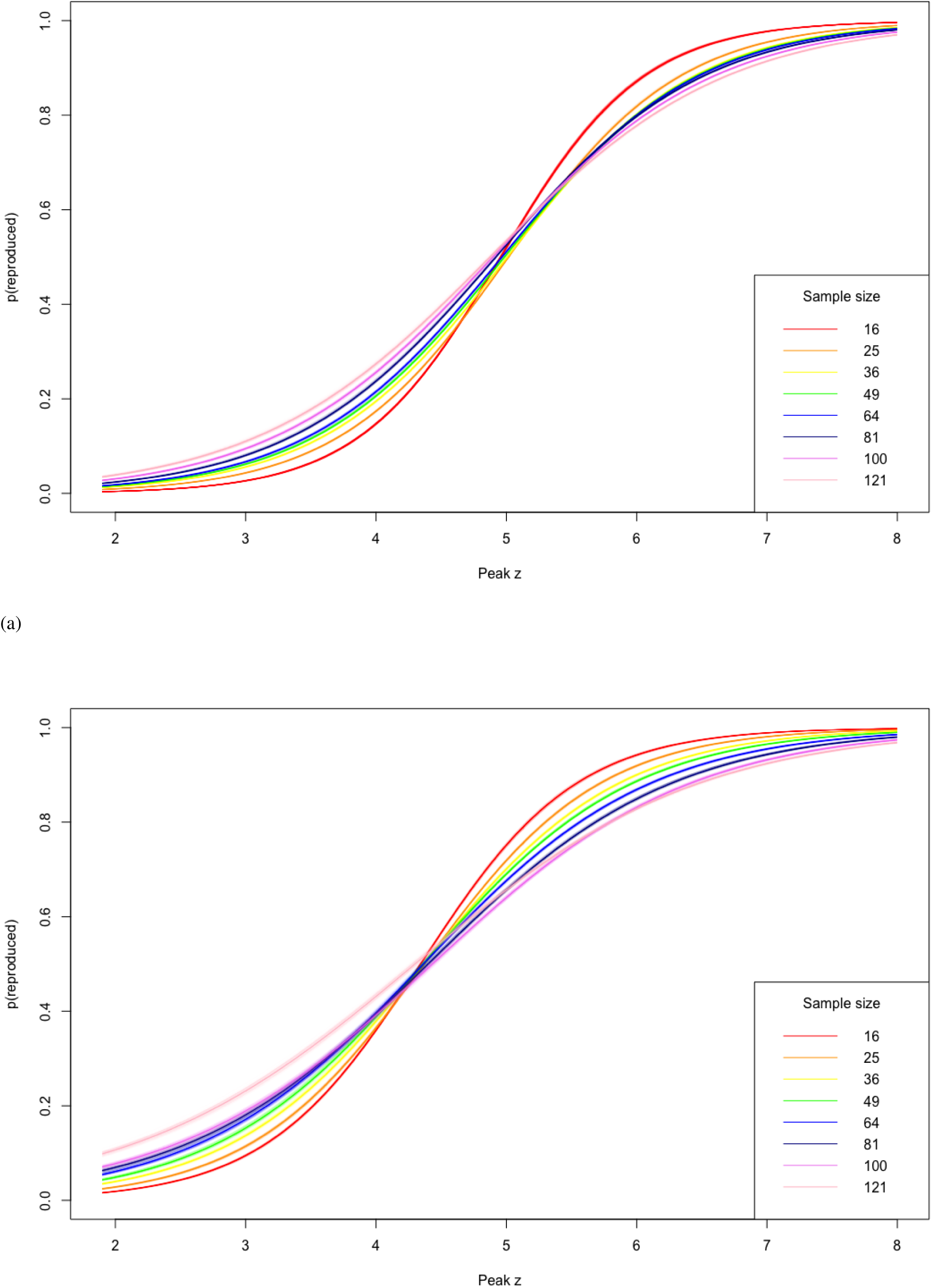
Replicability as a function of peak *z* value and sample size. (a) Conservative threshold; (b) Liberal threshold.

## Supplemental tables

**Table S1.** HCP tasks and contrasts.

| Task ID | Task name | Contrast number | Contrast description |
| --- | --- | --- | --- |
| E | EMOTION | 3 | Faces vs shapes |
| F | GAMBLING | 3 | Punish vs reward |
| G | LANGUAGE | 4 | Story vs math |
| H | MOTOR | 7 | Average motor vs fixation |
| I | RELATIONAL | 4 | Relation vs match |
| J | SOCIAL | 6 | Theory-of-mind vs random |
| K | WM | 11 | 2-back vs 0-back |

**Table S2.** Proportion suprathreshold in full-sample analyses for each task, using adaptive thresholds given in Table 1.

| Task | Proportion suprathreshold |  |
| --- | --- | --- |
|  | (liberal) | (conservative) |
| A | 0.115 | 0.025 |
| B | 0.260 | 0.131 |
| C | 0.429 | 0.227 |
| D | 0.133 | 0.044 |
| E | 0.344 | 0.191 |
| F | 0.149 | 0.040 |
| G | 0.669 | 0.498 |
| H | 0.444 | 0.273 |
| I | 0.269 | 0.116 |
| J | 0.432 | 0.271 |
| K | 0.408 | 0.250 |

**Table S3.** HCP subject IDs of all subjects included in HCP analyses.

|  |  |  |  |  |  |  |  |  |  |
| --- | --- | --- | --- | --- | --- | --- | --- | --- | --- |
| 100307 | 120212 | 141826 | 161327 | 181232 | 205725 | 298455 | 485757 | 654754 | 837560 |
| 100408 | 120515 | 142828 | 161630 | 182739 | 205826 | 303119 | 486759 | 665254 | 837964 |
| 101006 | 121618 | 143325 | 161731 | 183034 | 208024 | 303624 | 497865 | 672756 | 845458 |
| 101107 | 122317 | 144832 | 162026 | 185139 | 208226 | 304020 | 499566 | 673455 | 849971 |
| 101309 | 122620 | 145834 | 162228 | 185442 | 208327 | 307127 | 500222 | 677968 | 856766 |
| 101410 | 123117 | 146331 | 162329 | 186141 | 209834 | 308331 | 510326 | 679568 | 857263 |
| 101915 | 123420 | 146432 | 162733 | 187143 | 209935 | 310621 | 519950 | 680957 | 859671 |
| 102008 | 123925 | 147030 | 163129 | 187547 | 210011 | 316633 | 522434 | 683256 | 861456 |
| 102311 | 124220 | 147737 | 163331 | 187850 | 210415 | 329440 | 530635 | 687163 | 865363 |
| 102816 | 124422 | 148032 | 163432 | 188347 | 210617 | 334635 | 531536 | 690152 | 871762 |
| 103111 | 124826 | 148335 | 163836 | 189349 | 211215 | 339847 | 540436 | 695768 | 871964 |
| 103414 | 125525 | 148840 | 164030 | 189450 | 211316 | 351938 | 541943 | 702133 | 872764 |
| 103515 | 126325 | 148941 | 164131 | 190031 | 211417 | 352132 | 545345 | 704238 | 877269 |
| 103818 | 126628 | 149337 | 164939 | 191033 | 211720 | 352738 | 547046 | 705341 | 885975 |
| 104820 | 127630 | 149539 | 166438 | 191336 | 211922 | 355239 | 552544 | 709551 | 887373 |
| 105014 | 127933 | 149741 | 167036 | 191841 | 212116 | 356948 | 559053 | 713239 | 889579 |
| 105115 | 128127 | 150423 | 167743 | 192439 | 212217 | 361941 | 561242 | 715041 | 894673 |
| 105216 | 128632 | 150524 | 168139 | 192540 | 212318 | 365343 | 562446 | 729557 | 896778 |
| 106016 | 129028 | 150625 | 168341 | 192843 | 212419 | 366042 | 565452 | 732243 | 896879 |
| 106319 | 130013 | 150726 | 169444 | 193239 | 214019 | 366446 | 566454 | 734045 | 898176 |
| 106521 | 130316 | 151223 | 170934 | 194140 | 214221 | 371843 | 567052 | 742549 | 899885 |
| 107321 | 130922 | 151526 | 171633 | 194645 | 214423 | 377451 | 567961 | 748258 | 901038 |
| 107422 | 131217 | 151627 | 172029 | 194847 | 214726 | 380036 | 568963 | 749361 | 901139 |
| 108121 | 131722 | 151728 | 172130 | 195041 | 217126 | 382242 | 570243 | 751348 | 901442 |
| 108323 | 131924 | 152831 | 172332 | 195849 | 217429 | 385450 | 573249 | 753251 | 904044 |
| 108525 | 133019 | 153025 | 172534 | 196144 | 221319 | 386250 | 573451 | 756055 | 907656 |
| 108828 | 133625 | 153429 | 172938 | 196750 | 224022 | 390645 | 579665 | 759869 | 910241 |
| 109123 | 133827 | 153833 | 173334 | 197348 | 231928 | 395958 | 580044 | 761957 | 912447 |
| 109325 | 133928 | 154431 | 173435 | 197550 | 233326 | 397154 | 581349 | 765056 | 917255 |
| 110411 | 134324 | 154734 | 173536 | 198350 | 239944 | 397760 | 583858 | 770352 | 922854 |
| 111312 | 135225 | 154835 | 173940 | 198451 | 245333 | 397861 | 585862 | 771354 | 930449 |
| 111413 | 135528 | 154936 | 175035 | 198855 | 246133 | 412528 | 586460 | 779370 | 932554 |
| 111716 | 135932 | 155231 | 175439 | 199150 | 249947 | 414229 | 592455 | 782561 | 951457 |
| 113215 | 136227 | 155635 | 176542 | 199453 | 250427 | 415837 | 594156 | 784565 | 957974 |
| 113619 | 136833 | 156233 | 177645 | 199655 | 250932 | 422632 | 598568 | 786569 | 958976 |
| 113922 | 137027 | 156637 | 178142 | 199958 | 251833 | 433839 | 599469 | 788876 | 959574 |
| 114419 | 137128 | 157336 | 178748 | 200109 | 255639 | 436239 | 599671 | 789373 | 965367 |
| 114924 | 137633 | 157437 | 178849 | 200614 | 256540 | 436845 | 601127 | 792564 | 965771 |
| 115320 | 137936 | 158035 | 178950 | 201111 | 268850 | 441939 | 613538 | 792766 | 978578 |
| 116524 | 138231 | 158136 | 179346 | 201414 | 280739 | 445543 | 620434 | 802844 | 979984 |
| 117122 | 138534 | 158540 | 179548 | 201818 | 284646 | 448347 | 622236 | 814649 | 983773 |
| 117324 | 139233 | 159138 | 180129 | 203418 | 285345 | 465852 | 623844 | 816653 | 984472 |
| 118528 | 139637 | 159239 | 180432 | 204016 | 285446 | 473952 | 627549 | 826353 | 987983 |
| 118730 | 140117 | 159340 | 180836 | 204521 | 290136 | 475855 | 638049 | 826454 | 991267 |
| 118932 | 140824 | 159441 | 180937 | 205119 | 293748 | 479762 | 644044 | 833148 | 992774 |
| 119833 | 140925 | 160123 | 181131 | 205220 | 298051 | 480141 | 645551 | 833249 | 994273 |
| 120111 | 141422 | 160830 |  |  |  |  |  |  |  |

